# GABAergic LRP1 is a key link between obesity and memory function

**DOI:** 10.1101/2022.09.17.508390

**Authors:** Aaron Aykut Uner, Zhi-Shuai Hou, Ahmet Aydogan, Kellen C.C. Rodrigues, Jennie Young, Anthony Choi, Won-Mo Yang, Woojin S. Kim, Vincent Prevot, Barbara J. Caldarone, Bradley T. Hyman, Hyon Lee, Young-Bum Kim

**Author notes:** Present address: Key Laboratory of Mariculture (Ocean University of China), Ministry of Education (KLMME), Ocean University of China, Qingdao, China (Zhi-Shuai Hou). Present address: Department of Pathology, Faculty of Ceyhan Veterinary Medicine, Cukurova University, Adana, Turkey (Ahmet Aydogan). Present address: Department of Neurology, Gachon University Gil Medical Center, Incheon, Korea (Hyon Lee). Aaron Aykut Uner, Zhi-Shuai Hou and Ahmet Aydogan contributed equally to this work. Corresponding author: Division of Endocrinology, Diabetes and Metabolism, Beth Israel Deaconess Medical Center, 330 Brookline Avenue, Boston, MA, 02215 USA. (Y.B. Kim). Declarations of interest: None.

## Abstract

**Objective:** Low-density lipoprotein receptor-related protein-1 (LRP1) regulates energy homeostasis, blood-brain barrier integrity, and metabolic signaling in the brain. Loss of LRP1 from inhibitory gamma-aminobutyric acid (GABA)ergic neurons causes severe obesity in mice. Its dysfunction has been associated with cognitive decline, dementia, and Alzheimer’s disease. However, the impact of LRP1 in inhibitory neurons on memory function and cognition in the context of obesity is poorly understood.

**Methods:** Mice lacking LRP1 in GABAergic neurons (*Vgat-Cre; LRP1^loxP/loxP^*) are subjected to conduct behavioral tests of locomotor activity and motor coordination, short/long-term and spatial memory, and fear learning/memory. We evaluated the relationships between behavior and metabolic risk factors.

**Results:** Deletion of LRP1 in GABAergic neurons caused a significant impairment in memory function. In the spatial Y-maze test, *Vgat-Cre; LRP1^loxP/loxP^* mice exhibited decreased travel distance and duration in the novel arm compared with controls (*LRP1^loxP/loxP^* mice). In addition, GABAergic neuron-specific LRP1-deficient mice had a diminished capacity for performing learning and memory tasks during the water T-maze test. Moreover, reduced freezing time was observed in these mice when the contextual and cued fear conditioning tests were conducted. These effects were accompanied by increased neuronal necrosis and neuroinflammation in the hippocampus. Importantly, the distance and duration in the novel arm and the performance of the reversal water T-maze test negatively correlated with metabolic risk parameters, including body weight, serum leptin, insulin, and apolipoprotein J.

**Conclusions:** Our findings demonstrate that LRP1 from GABAergic neurons is important in normal memory function. Metabolically, obesity caused by GABAergic LRP1 deletion negatively regulates memory and cognitive function. Thus, LRP1 in GABAergic neurons may play a crucial role in maintaining normal excitatory/inhibitory balance and impacts memory function, reinforcing the potential importance of LRP1 in neural system integrity.

## 1. INTRODUCTION

Emerging data from human and animal studies suggest that obesity is linked to cognitive decline, brain atrophy, reduced white matter volume and integrity of the blood-brain barrier, and an elevated risk of late-onset Alzheimer’s disease [1-6]. Chronic disturbances in glucose homeostasis, impaired insulin signaling, and hypometabolism are highly correlated with cognitive impairment and Alzheimer’s disease pathology [7-13]. We and others demonstrated that genetic disruption of low-density lipoprotein receptor-related protein 1 (LRP1) in the central nervous system or selectively in inhibitory gamma-aminobutyric acid (GABA)ergic neurons results in increased food intake, decreased energy expenditure, and metabolic changes leading to obesity [14,15]. LRP1, a member of the low-density lipoprotein (LDL) receptor family, serves as a multi-ligand cell surface receptor and is broadly expressed in neurons [14,16]. In addition to its critical role in metabolism, LRP1 is also a major regulator of neurotransmission, synaptic plasticity, and amyloid-β (Aβ) clearance [17-20]. Aberrant LRP1 expression has been associated with the development of cognitive dysfunction and Alzheimer’s disease-related dementia [21]. In particular, LRP1 levels were found to be markedly reduced in the brain of patients with Alzheimer’s disease and decreased LRP1 levels correlated with increased Alzheimer’s disease susceptibility [22]. In line with this, evidence revealed that the deletion of LRP1 in endothelial cells of the brain led to a significant deficit in spatial learning and memory [23]. However, the impact of LRP1 selectively in GABAergic neurons on cognition is unknown.

In the current study, we investigated the role of LRP1 in GABAergic neurons in memory and cognitive function as well as the relationship between obesity-induced metabolic changes and behavioral outcomes.

## 2. MATERIALS AND METHODS

### 2.1. Animal care

All animal care and experimental procedures were conducted in accordance with the National Institute of Health’s Guide for the Care and the Use of Laboratory Animals and approved by the Institutional Animal Care and Use Committee of Beth Israel Deaconess Medical Center and Harvard Medical School (HMS). Mice were allowed access to a standard chow diet [LabDiet, 5008 (5058 at HMS) Formulab diet, Irradiated] and water (reverse osmosis water) was provided *ad libitum.* They were housed at 22–24 °C with a 12-h light-dark cycle with the light cycle starting from 6:00 am to 6:00 pm (7:00 am – 7:00 pm at HMS). Mice were single-housed for behavioral studies.

### 2.2. Generation of Vgat-Cre; LRP1^loxP/loxP^ mice

Mice bearing a *loxP*-flanked *LRP1* allele (*LRP1^loxP/loxP^*) were purchased from The Jackson Lab (Stock No: 012604, Bar Harbor, ME). Mice lacking LRP1 in the vesicular GABA transporter (Vgat)-expressing neurons (*Vgat-Cre; LRP1^loxP/loxP^*) were generated by mating *LRP1^loxP/loxP^* mice with *Vgat-cre* mice (gift from Dr. Brad Lowell, Beth Israel Deaconess Medical Center, Boston, MA). These mice were derived from 129 ES cells in C57BL/6 embryos. All mice studied were mixed background with 129 and C57BL/6. LRP1 expression was reduced in the arcuate nucleus, paraventricular hypothalamic nucleus, and dorsomedial hypothalamic nucleus of *Vgat-Cre; LRP1^loxP/loxP^* mice where *Vgat* is expressed [14]. *Vgat-Cre; LRP1^loxP/loxP^* and *LRP1^loxP/loxP^* littermates were used for analyses. Only healthy mice showing normal behavior, development and constant physical activity (even though they are obese) were used. Mice were excluded from the experiments if they show abnormal development such as hydrocephalus.

### 2.3. Measurements of A**β** and metabolic parameters

Mice were weighed from 4 weeks of birth and weekly thereafter. Epidydimal fat was harvested and weighed at the end of the experiment in male mice at 32 weeks of age. Blood was collected from overnight fasted mice via the tail. Blood glucose was measured using a OneTouch Ultra glucose meter (LifeScan, Inc., Milpitas, CA). Serum levels of insulin (90080, Crystal Chem, Chicago, IL), leptin (90030, Crystal Chem, Chicago, IL), apolipoprotein J (ApoJ) (MCLU00, R&D Systems, Minneapolis, MN), Aβ_40_ (KMB3481, Thermo Fisher Scientific Inc., Waltham, MA), Aβ_42_ (KMB3441, Thermo Fisher Scientific Inc., Waltham, MA) and free fatty acids (FFA) (ab65341, Abcam, Waltham, MA) were measured by enzyme linked immunosorbent assay (ELISA). The homeostatic model assessment for insulin resistance (HOMA-IR) was calculated as [fasting glucose (mmol/L) × fasting insulin (μIU/mL)]/22.5 with a slight modification in insulin conversion as has recently been recommended [24,25].

### 2.4. Immunoblotting analysis

Tissue lysates were resolved by sodium dodecyl sulfate–polyacrylamide gel electrophoresis and transferred to nitrocellulose membranes (GE Healthcare Life Sciences, Pittsburgh, PA).

The membranes were incubated with polyclonal antibodies against presenilin-1 (PSEN1) (sc-365450, Santa Cruz Biotechnology Inc., Dallas, TX) and amyloid precursor protein (APP) (A8717, Sigma-Aldrich, St. Louis, MO) or monoclonal antibody against β-actin (A2228, Sigma-Aldrich, Louis, MO). The membranes were washed with Tris-buffered saline (TBS) containing 0.1% Tween 20 for 30 min, incubated with horseradish peroxidase secondary antibodies against mouse (7076, Cell Signaling Technology, Danvers, MA) or rabbit (7074, Cell Signaling Technology) for 1 h, and washed with TBS containing 0.1% Tween 20 for 30 min. The bands were visualized with enhanced chemiluminescence and quantified by an ImageJ program (v1.52a, NIH). All protein levels were normalized by β-actin levels.

### 2.5. Locomotor activity

Mice were placed in the center of a square Plexiglas box (27 cm × 27 cm × 20.3 cm) containing infrared arrays (Med Associates, St Albans, VT, USA) and allowed to freely explore the environment for 1 h. The total activity (cm) was recorded using specialized software (Activity Monitor, Version 5.9).

### 2.6. Spatial Novelty Y Maze

The Spatial Novelty Y Maze is a two-trial short-term memory test that measures a mouse’s preference to explore a novel environment. In the first forced trial, the mouse is placed in the “start” arm and is allowed to explore both the “start” and “familiar” arm. After a short delay, the mouse is given the free choice to explore the “familiar” or “novel’ arm. If the mouse remembers the familiar arm, it should choose to explore the “novel” more than the “familiar” arm. Testing was conducted in a clear acrylic Y maze with three arms (one start arm and two test arms, all ∼31 cm in length). The testing room had distinct visual cues and the maze had a removable partition to block the appropriate arm. The blocked arm was randomized and balanced for each genotype. The test consisted of a forced choice trial followed by a free-choice trial. For the forced choice trial, the start arm and one test arm were open with access to the second test arm blocked by the partition. Individual subjects were placed in the start arm and allowed to explore the open test arm for 3 min, after which they were removed from the maze, and placed in a holding cage as the maze was cleaned. The partition was removed and the mice were then placed back into the Y maze for the free choice trial and allowed to explore both the open and test arms for 3 min. The delay between the forced choice and free choice trials was 2 min. Animal behavior was video recorded during both trials and the time spent in the previously accessible arm (i.e. familiar arm) and the previously blocked arm (i.e. novel arm) is determined during the free choice trial using Ethovision software. For each subject the % time and distance exploring the novel arm during the free choice trial is calculated using the formula: % novel = novel / (novel + familiar) x 100.

### 2.7. Rotarod test

Motor coordination was assessed using a rotarod test (Ugo Basile, Gemonio, VA, Italy). Mice were first given a 5 min habituation trial in which the mice were placed on the apparatus that was rotating at a constant speed of 4 rpm. If mice fell off during this acclimation period, they were placed back on the rod. Mice were then given two test trials, in which the rod accelerated at a 4 – 40 rpm over a course of 3 min. The maximum cutoff time was 5 min. When mice fell off the rod, the latency to fall was recorded. Mice were given a 3 h interval between trials. The average of the two trials was calculated.

### 2.8. Water T-maze

A custom made Plexiglas plus maze testing apparatus (each arm 36 cm length, 10 cm width, 21 cm height) (Plastic Craft, West Nyack, NY) with four arms (North, South, East and West) was used to evaluate spatial memory. One arm was blocked off with a divider so that the mouse could choose one of the east and west arms to escape. An escape platform was placed on the east side of the maze ∼ 1 cm below the surface water. The maze was filled with 23-26 °C water and white non-toxic paint to ensure that the mouse could not see the escape platform. After a 10 min acclimation period in the testing room, the mouse was placed into the appropriate arm facing the wall and was allowed to swim until it found the escape platform. The start position was alternated (north and south) in a semi randomized order. If the mouse was not able to find the escape platform within 1 min, the experimenter guided the mouse to the platform. The mouse was allowed to stay on the platform for 10 sec before being placed back into its holding cage. Mice were tested for 5 days (10 trials/day) and correct and incorrect choices to find the escape platform were recorded. Once both knock-out and control mice reached the acquisition criteria, which was a group mean of 80% or more correct responses, the escape platform was moved to the opposite side (west) and reversal learning was conducted for 5 days.

### 2.9. Contextual and cued fear conditioning tasks (CCFC)

Mice were placed in Plexiglas conditioning chambers (Med Associates, St. Albans, VT) with speakers (Med Associates Sound generator ENV-230, St. Albans, VT) and steel bars spaced 1 cm apart for electric shocks (0.5 mA). On the first day (training phase), after a 2 min acclimation period, the mice received two tones (30 sec) and electric shocks (2 sec) within the last two seconds of the tone with a 2 min inter-trial interval. On day 2, the mice were placed in the chamber, but did not receive a tone and electric shock. Freezing behavior was recorded for 5 min in the conditioning context and analyzed by the Ethovision Software (EthoVision XT 14, Noldus, Wageningen, the Netherlands). After all mice were tested for contextual fear conditioning, the context was changed by inserting a white Plexiglas floor over the shock grid and white Plexiglas to cover the wall. Freezing was measured for 3 min in this altered context and for 3 min during the presentation of the tone.

### 2.10. Histopathology

Mice were transcardially perfused with phosphate-buffered saline (PBS) followed by a 10% formalin solution. Brain samples obtained from the mice were further fixed in a 10% neutral formalin solution overnight and cut sagittally (lateral from ∼ 0.24 mm to 2.04 mm). They were then processed using graded alcohols and xylene, embedded in paraffin, sectioned at 5 μm, stained with hematoxylin and eosin (H&E) (MHS32, Sigma-Aldrich, Louis, MO), and evaluated under a light microscope (Olympus BH-2, Shinjuku, Tokyo, Japan and Nikon Eclipse E 600, Tokyo, Japan) [26]. Necrotic neurons with perineural satellitosis as well as microglial cells in the hippocampus were counted manually under a light microscope with 40× magnification in 15 different areas from five serial sections.

### 2.11. Immunohistochemical staining

Immunohistochemistry (IHC) was performed on 5-μm paraffin sections using the standard avidin–biotin–peroxidase complex (ABC) method according to the manufacturer’s recommendations (Mouse and Rabbit Specific HRP/DAB (ABC) Detection IHC kit (ab64264), Abcam, Waltham, MA). Sections were subjected to routine deparaffinization in xylene and then rehydration in ethanol (100%, 80%, and 50%). Inhibition of endogenous peroxidase activity and blocking of non-specific binding were performed according to the kit’s instructions. Antigen retrieval for all markers was done by microwaving the sections in citrate buffer (pH = 6.0) at a sub-boiling temperature for 10 min. After a second protein blocking, brain sections were incubated with primary antibodies overnight at 4 °C. The primary antibodies used were anti-glial fibrillary acidic protein (GFAP) antibody (dilution; 1:100, Sigma-Aldrich (G6171), St. Louis, MO), anti-ionized calcium binding adaptor molecule 1 (IBA1) antibody (dilution; 1:100, Wako (019-19741), Richmond, VA), anti-glutamic acid decarboxylase 67 (GAD67) antibody (clone 1G10.2, dilution; 1:100, Sigma-Aldrich (MAB5406)) and anti-N-methyl-D-aspartate receptor 1 (NMDAR1) antibody (dilution; 1:100, Abcam (ab193310)). The brown color of immunopositivity in brain sections was developed with 3,3’-diaminobenzidine tetrahydrochloride (DAB-H2O2) substrate in PBS for 5 min. Slides were counterstained with Mayer’s hematoxylin (Sigma-Aldrich), dehydrated, and mounted with Entellan (Sigma-Aldrich). On each slide, immunopositive reactions were detected by the presence of brown cytoplasmic staining. Immunopositive cells were counted as described in the histopathology section.

### 2.12. Double immunofluorescent staining

To determine if knock down of LRP1 from GABAergic neurons changes the PSEN1 expression in GABAergic neurons and LRP1 expressions in astrocytes and microglial cells, we performed double immunofluorescent staining. Mice were transcardially perfused as described in the histopathology section. Brains were removed, incubated in 10% formalin and then 20% sucrose solution (overnight each), cut coronally on a microtome into sections (25 µm), and collected in four series. Five sections between - 1.46 mm and - 2.80 mm from the Bregma were picked, washed with PBS, permeabilized in 0.25% Triton X-100 (15 min at room temperature), blocked in 10% normal donkey serum (1 h at room temperature), and then incubated overnight in blocking solution containing primary antibody [Lrp1 (rabbit) dilution; 1:500, Abcam (ab92544); Lrp1 (mouse) dilution; 1:100, Invitrogen (37-3800); GAD67 (clone 1G10.2) dilution; 1:200, Sigma-Aldrich (MAB5406); GFAP dilution; 1:200, Sigma-Aldrich (G6171); IBA1 dilution; 1:200, Wako (019-19741) and PSEN1 dilution; 1:50, Thermo Fisher Scientific (PA5-119872)] at room temperature. The next day, the sections were washed and then incubated in secondary antibodies [Alexa fluor (AF) 488, AF 594 and AF 647 dilution; 1:500]. After washing, the sections were mounted onto slides and images were taken with a slide scanner microscope (Olympus VS120). Double immunofluorescent positive cells (LRP1 vs GAD67, LRP1 vs IBA-1, LRP1 vs GFAP and PSEN1 vs GAD67) (100 cells in each section) in the hippocampus were counted manually in the five serial sections.

### 2.13. Statistical analysis

A power analysis was conducted using G Power (Version 3.1.9.2, Germany) to estimate the sample size. Probability of α and β errors was set 0.05 and 0.20, respectively. Effect size was calculated according to software’s instructions or preliminary data if any. The data were checked for the normality of distribution by the Shapiro-Wilk test. Logarithmic transformation or non-parametric tests were done if the data were not distributed normally. The student t-test or Mann-Whitney U test were conducted to compare two groups. The paired t-test or Wilcoxon test was used when comparing the data within groups. Two-way analysis of variance (ANOVA) was done when two independent factors [e.g. time-group or arm (familiar/novel)-group] might have an effect. *Post hoc* multiple comparisons were carried out using Sidak test in GraphPad Prism (GraphPad Prism, Version 8.0.1, La Jolla, CA) or general linear procedures in SPSS (extra coding were applied on the syntax menu) (SPSS 26.0 software for Windows; SPSS Inc., Chicago, IL). Body weight was used as covariate when appropriate. Correlations were done using Pearson’s or Spearman correlation analyses depending on the data distribution. *P* ≤ 0.05 was considered significant. The results were presented as mean ± standard error of mean (s.e.m.).

## 3. RESULTS

### 3.1. Deletion of LRP1 in GABAergic neurons causes severe obesity

Double immunofluorescent staining revealed that LRP1-containing neurons in GABAergic neurons are significantly decreased ∼45% in *Vgat-Cre; LRP1^loxP/loxP^* mice compared with *LRP1^loxP/loxP^* mice (Figure 1A). Since LRP1 deletion from GABAergic neurons may cause compensatory changes that can affect overall memory function, we measured LRP1 levels in microglial cells and astrocytes as well as PSEN1 levels in GABAergic neurons. We found that LRP1 (∼ 6%) and PSEN1 (∼ 4%) slightly decreased in microglial cells (Figure 1B) and GABAergic neurons (Figure 1C), respectively, while LRP1 expression in astrocytes did not change (Figure 1D). We observed that mice lacking LRP1 in GABAergic neurons were severely obese (Figure 1E) with increases in epididymal fat (Figure 1F). Serum levels of leptin (Figure 1G) and insulin (Figure 1J) were also increased in *Vgat-Cre; LRP1^loxP/loxP^* mice compared with *LRP1^loxP/loxP^* mice at 32 weeks of age. However, neither serum FFA nor blood glucose levels were altered in *Vgat-Cre; LRP1^loxP/loxP^* mice (Figure 1H-I). *Vgat-Cre; LRP1^loxP/loxP^* mice were insulin resistant as demonstrated by increases in the HOMA-IR index (Figure 1K). These results suggest that LRP1 activity in GABAergic neurons is required to regulate bodyweight and metabolic homeostasis and are consistent with our previous report showing the deletion of LRP1 from GABAergic neurons leads to severe obesity by increasing food intake and decreasing energy expenditure and locomotor activity [14].

**Figure 1:**
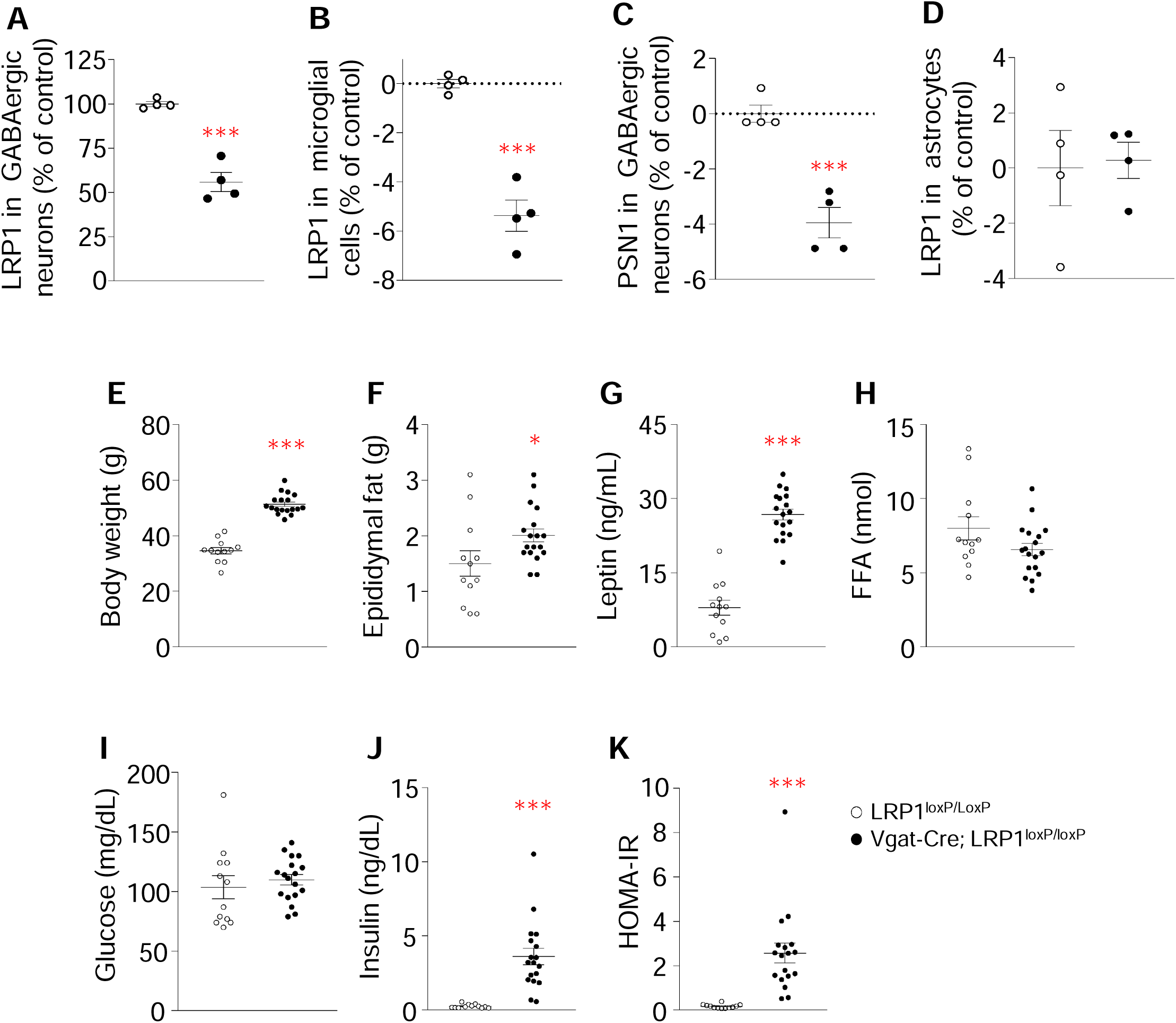
Selective deletion of LRP1 in GABAergic neurons causes metabolic disorders. Changes in LRP1 in (**A**) GABAergic neurons, (**B**) microglial cells and (**D**) astrocytes and PSEN1 levels in (**C**) GABAergic neurons. (**E**) Body weight, (**F**) epididymal fat, (**G**) serum leptin, (**H**) serum FFA, (**I**) blood glucose, (**J**) serum insulin and (**K**) HOMA-IR of *LRP1^loxP/loxP^* and *Vgat-Cre; LRP1^loxP/loxP^* male mice. Body weight, epididymal fat, glucose, and serum parameters were measured from overnight fasted mice at 32 weeks of age. *n* = 12 for control, *n* = 18 for *Vgat-Cre; LRP1^loxP/loxP^*. APP and PSEN1 protein levels were assessed by immunoblotting. LRP1 levels in GABAergic neurons, microglial cells and astrocytes and PSEN1 levels in GABAergic neurons were determined by double immunofluorescent staining from overnight fasted mice at 28 weeks of age (*n* = 4/group). All graphs show means ± SEM. \**P* <0.05, \*\*\**P* <0.001 vs. *LRP1^loxP/loxP^* by two-sided Student’s t-test or Mann-Whitney U test.

### 3.2 Mice lacking LRP1 in GABAergic neurons have increased serum ApoJ levels

ApoJ has been implicated in the pathogenesis of Alzheimer’s disease [27-30]. Given that LRP1 is a potential receptor for ApoJ [31,32] and serum ApoJ levels are elevated in humans with obesity and type 2 diabetes as well as diet-induced obese mice [33-35], we measured ApoJ levels in the serum and tissues of *Vgat-Cre; LRP1^loxP/loxP^* mice. We found that serum ApoJ levels (Supplementary Figure 1A) were significantly elevated in *Vgat-Cre; LRP1^loxP/loxP^* mice compared to *LRP1^loxP/loxP^* mice, whereas ApoJ levels in the liver, brain, hippocampus, and cerebral cortex remained unaltered (Supplementary Figure 1B–E). Furthermore, there was a strong correlation between serum ApoJ levels and body weight, serum insulin, and leptin levels (Supplementary Figure 1F–H). These findings suggest that increases in serum ApoJ levels are associated with LRP1-mediated metabolic dysfunction.

### 3.3. LRP1 in GABAergic neurons regulates motor function and spatial recognition memory

Locomotor activity and rotarod tests were assessed to determine whether GABAergic deletion of LRP1 alters motor activity and coordination. *Vgat-Cre; LRP1^loxP/loxP^* mice displayed a significant decrease in the total distance travelled over 1 h (Figure 2A) and in 5 min intervals (Figure 2B) compared with *LRP1^loxP/loxP^* mice. These effects were associated with reduced rotarod performance in *Vgat-Cre; LRP1^loxP/loxP^* mice (Figure 2C). When the body weights of mice were used as covariates, we found that the total distance (Figure 2A) and the distance travelled in 5 min intervals (Figure 2B) were no longer significantly different between groups while the values of motor coordination remained significant (Figure 2C). Mice were subjected to the Y-maze test to determine whether LRP1 activity in GABAergic neurons is involved in short-term memory. The distance traveled and time spent in the familiar arm were similar between *LRP1^loxP/loxP^* and *Vgat-Cre; LRP1^loxP/loxP^* mice (left panels of Figure 2D and F), and the distance traveled and time spent in the novel arm were increased compared with the familiar arm for both groups (right panels of Figure 2D and F), indicating that both groups are capable of learning and memory. However, mice lacking LRP1 in GABAergic neurons visited the novel arm less frequently (Figure 2D left panel and E) and spent less time there (Figure 2F left panel and G) than *LRP1^loxP/loxP^* mice. These observations demonstrate that LRP1 in GABAergic neurons may play a role in regulating motor function and spatial short-term memory.

**Figure 2:**
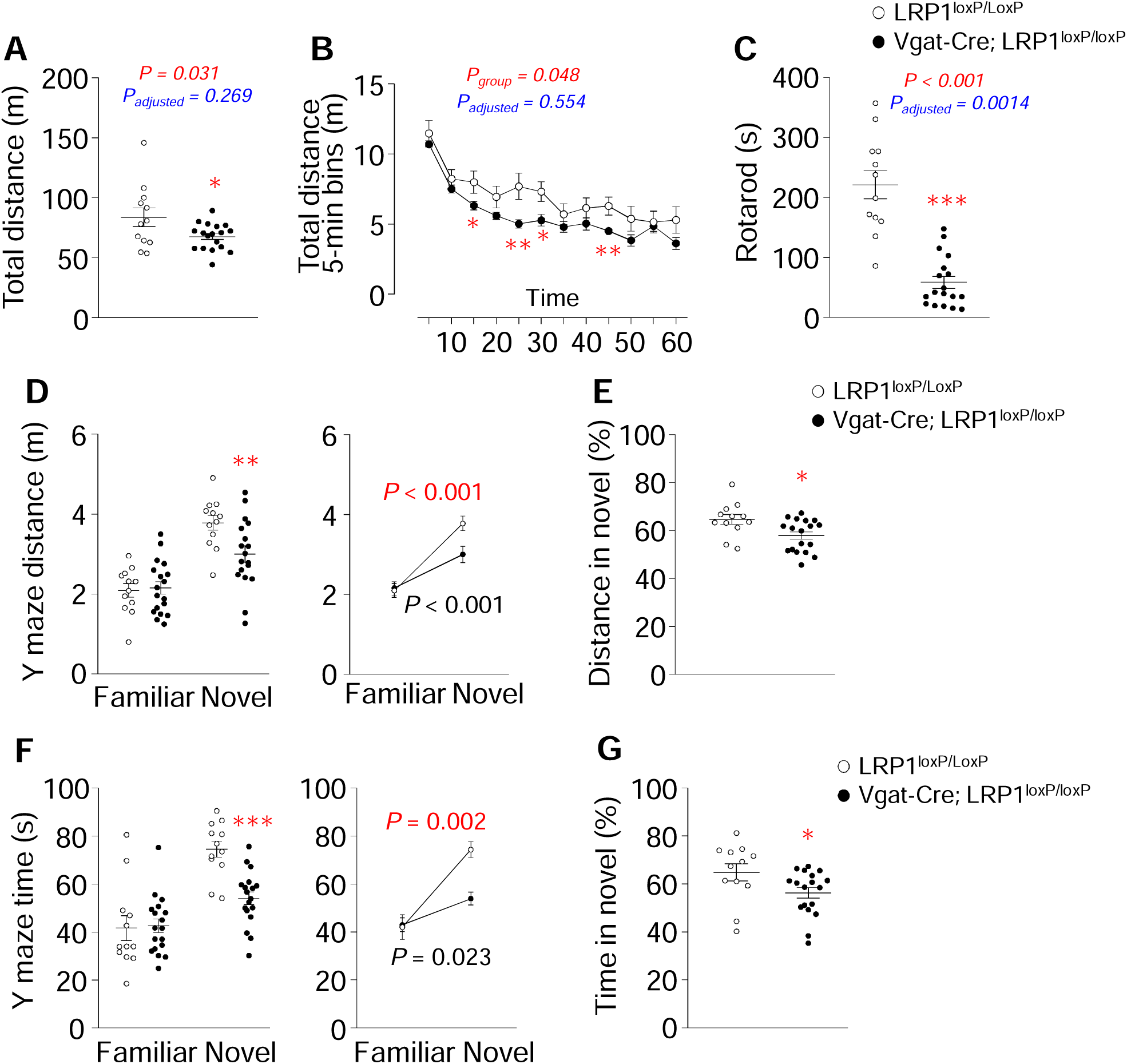
GABAergic neuron-specific deletion of LRP1 alters locomotor activity, motor coordination and spatial recognition memory. (**A**) Total distance travelled in 1 h on the locomotion test, (**B**) distance travelled in 5 min intervals on the locomotion test, (**C**) latency to fall on the rotarod test, (**D**) total distance travelled in novel and familiar arms on the spatial Y-maze test (left panel), and changes in total distance travelled from the familiar to novel arm (right panel), (**E**) percentage of distance travelled in the novel arm of spatial Y-maze, (**F**) time spent in the novel and familiar arms on the spatial Y-maze test (left panel), and changes in time spent from the familiar to novel arm (right panel), and (**G**) percentage of time spent in the novel arm of the spatial Y-maze. *n* = 12 for control, *n* = 18 for *LRP1^loxP/loxP^* (26 weeks old). All graphs show means ± SEM. \**P* <0.05, \*\**P* <0.01, \*\*\**P* <0.001 vs. *LRP1^loxP/loxP^* by two-sided Student’s t-test. Data in right panels of D and F were evaluated by paired t-test. The adjusted *P* values in A, B and C were calculated by the covariance analysis (Body weight was used as covariate).

### 3.4. LRP1 in GABAergic neurons is involved in cognitive function

We tested mice in the water T-maze and CCFC to further determine the impact of LRP1 activity on cognitive function. In the water T-maze test, the average percentage of correct responses to find the hidden escape platform on the first day of acquisition was lower in *Vgat-Cre; LRP1^loxP/loxP^* mice compared to controls, but no differences were observed on day 2 to 5 (Figure 3A). Similar results were found when the escape platform was placed on the opposite side (reversal) (Figure 3B). In the CCFC test, *Vgat-Cre; LRP1^loxP/loxP^* mice displayed a higher baseline freezing time on the training day than LRP1*^loxP/loxP^* mice. We therefore adjusted the results [including the training day, (Figure 3C)] according to the baseline values of the training day. The adjusted data revealed that on the training day, all mice increased their freezing time slightly or significantly on tone 2 and shock 2 compared to tone 1 and shock 1 (Figure 3C), indicating a recognition of the neutral (tone) and aversive (shock) stimuli. However, *Vgat-Cre; LRP1^loxP/loxP^* mice exhibited less freezing time compared to the control mice on the second day in CCFC tests (Figure 3C). Together, these findings suggest that the loss of LRP1 from GABAergic neurons leads to impairments in learning and long-term memory.

**Figure 3:**
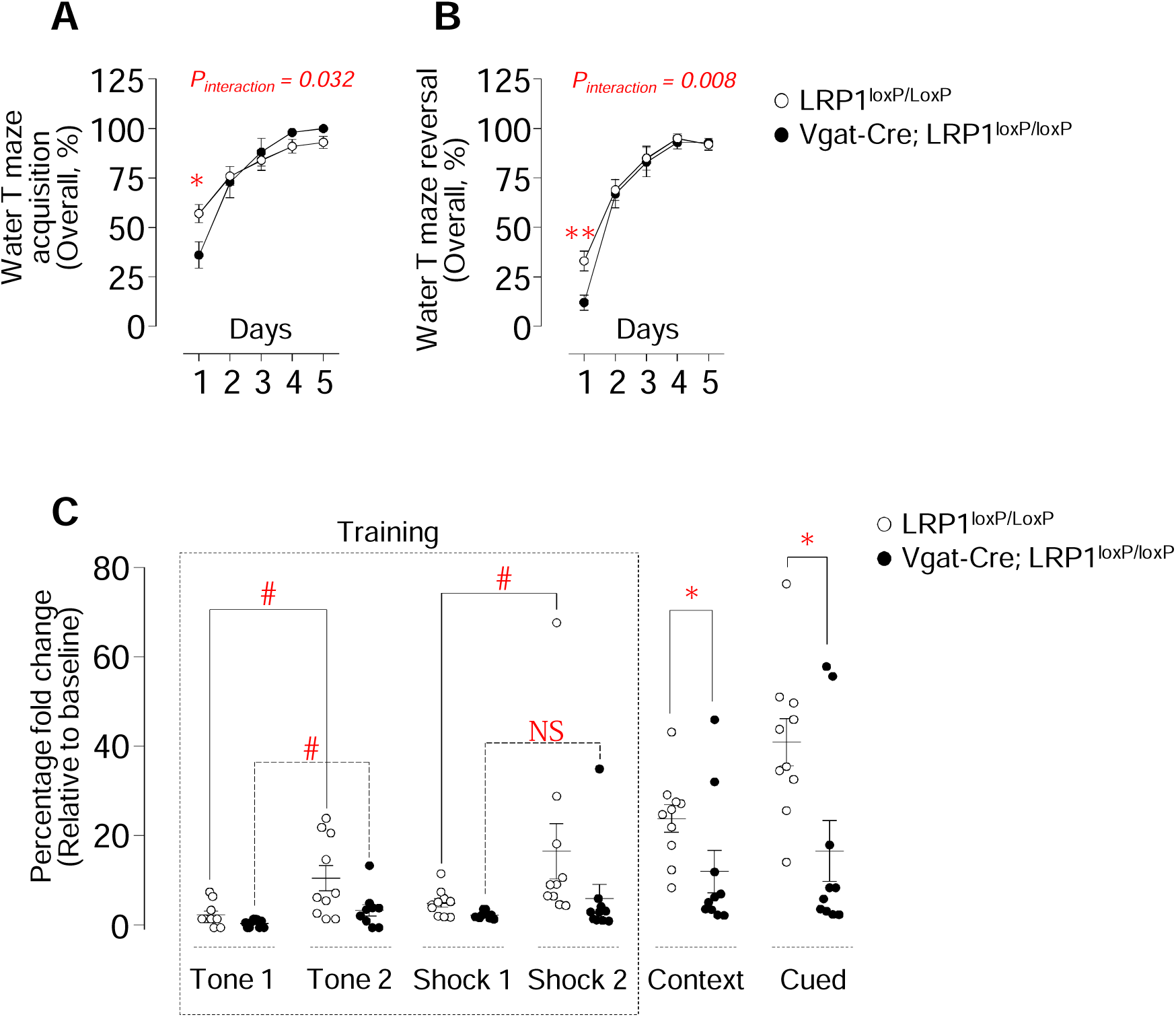
Loss of LRP1 in GABAergic neurons leads to memory dysfunction. Percentage correct response (10 trials/day) of *LRP1^loxP/loxP^* (*n* = 10) and *Vgat-Cre; LRP1^loxP/loxP^* (*n* = 10) male mice (28-29 weeks old) on the (**A**) acquisition and (**B**) reversal water T-maze. \**P* <0.05, \*\**P* <0.01 by two-way ANOVA (*post hoc* tests were done using GLM procedures on SPSS). (**C**) Percentage fold change (relative to baseline) in freezing time of *LRP1^loxP/loxP^* (*n* = 10) and *Vgat-Cre; LRP1^loxP/loxP^* (*n* = 10) male mice (31 weeks old) during the fear conditioning test. All graphs show means ± SEM. \**P* <0.05 by two-sided Student’s t-test. ^#^*P* <0.05 by paired t-test or Wilcoxon test. NS: Not significant.

### 3.5. Perturbed metabolic profiles are associated with cognitive dysfunction in mice lacking GABAergic

We performed correlation analyses between behavioral and metabolic parameters to determine the relationship between cognitive function and metabolic profiles. Total travel distance and motor coordination negatively correlated with body weight (Figure 4A–B). Total travel distance and duration in the novel arm, as measured by the Y-maze test, negatively correlated with metabolic parameters, including body weight (Figure 4C–D), serum insulin (Figure 4E–F), leptin (Figure 4G–H), and ApoJ (Figure 4I). These results suggest a potential relationship between obesity and dysregulated locomotor activity, motor coordination, and spatial recognition memory.

**Figure 4:**
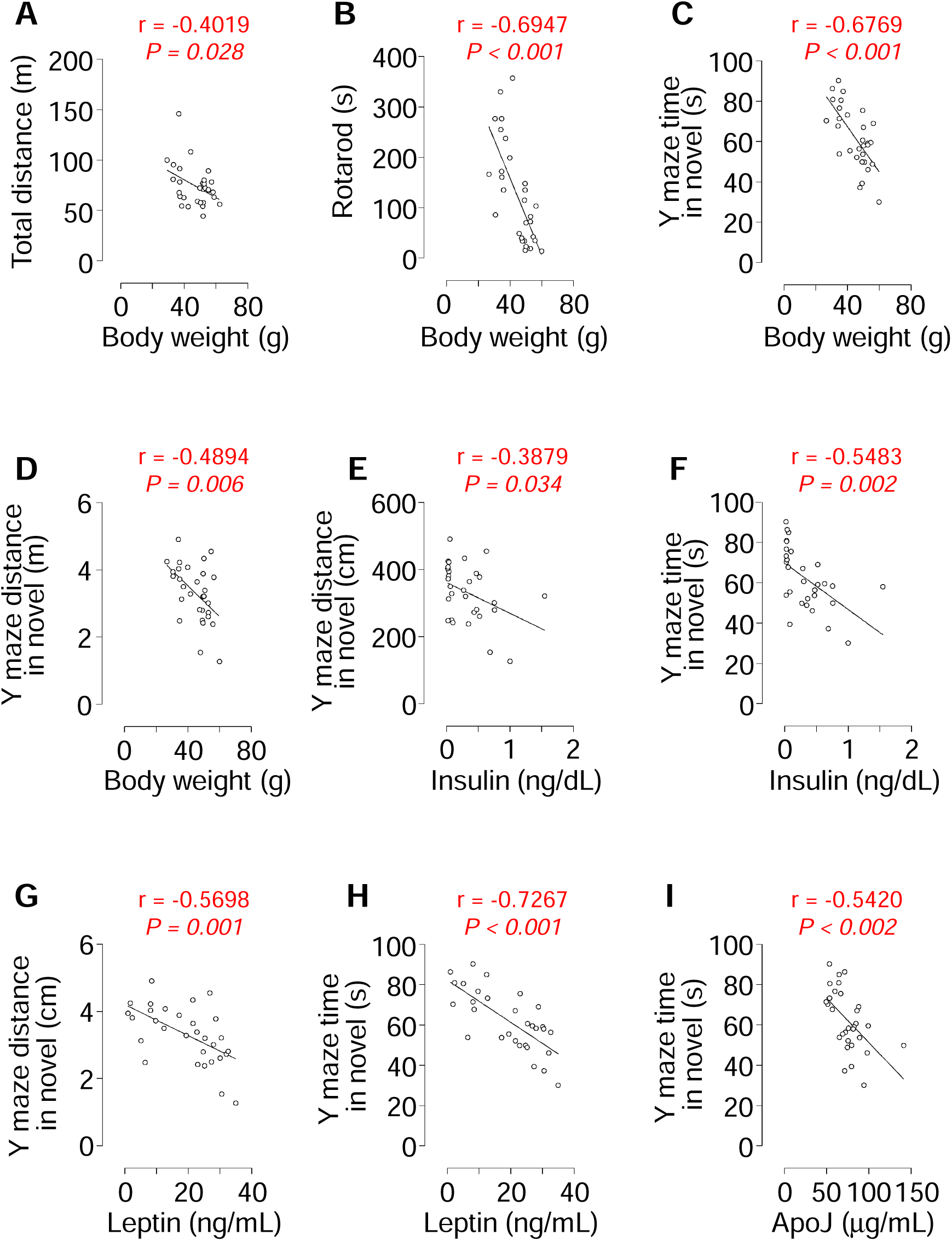
Relationship of motor function, motor coordination and spatial recognition memory with obesity-linked metabolic parameters. Correlation of body weight with (**A**) total distance travelled on the locomotion test, (**B**) latency to fall on the rotarod test, (**C**) total distance travelled and (**D**) time spent in the novel arm on the spatial Y-maze test. Correlation of serum insulin with (**E**) total distance travelled and (**F**) time spent in the novel arm on the spatial Y-maze test. Correlation of serum leptin with (**G**) total distance travelled and (**H**) time spent in the novel arm on the spatial Y-maze test. (**I**) Correlation of serum ApoJ with time spent in the novel arm on the spatial Y-maze test. The *P* values were obtained by Pearson correlation analysis and r values indicate Pearson correlation coefficient.

Similar to the Y-maze correlation results, body weight (Figure 5B), and the serum levels of leptin (Figure 5D), insulin (Figure 5F) and ApoJ (Figure 5H) were found to be negatively correlated with the data from the reversal water T-maze test. Serum ApoJ levels (Figure 5G) also negatively correlated with the water T-maze acquisition, whereas body weight (Figure 5A), serum leptin (Figure 5C), and insulin (Figure 5E) did not correlate with the water T-maze acquisition. These data indicate that perturbed metabolic outcomes caused by LRP1 deletion in GABAergic neurons are associated with impaired cognitive function.

**Figure 5:**
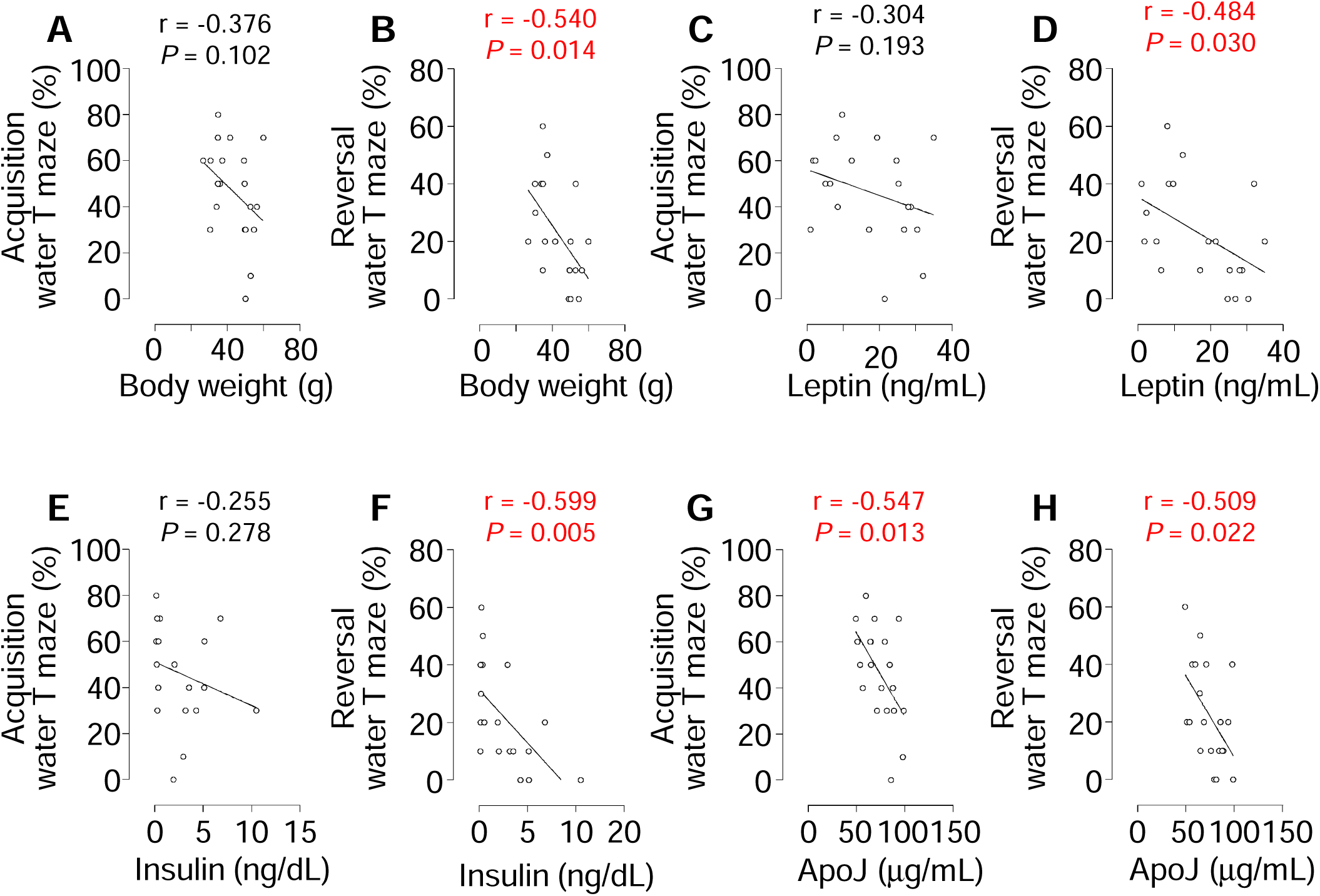
Relationship between metabolic parameters and cognitive function. Correlation of body weight, serum leptin, serum insulin, and serum ApoJ with the data from acquisition (**A, C, E, G**) and reversal (**B, D, F, H**) water T-maze on day 1. The *P* values were obtained by Pearson correlation analysis and r values indicate Pearson correlation coefficient.

### 3.6. The impact of LRP1 on A**β** accumulation

Since LRP1 plays important role in Aβ clearance [19], we measured Aβ levels in the serum, brain, and liver to understand the impact of GABAergic neuron-specific LRP1-deficiency on Aβ production. The levels of Aβ_42_ trended higher in the serum and brain of *Vgat-Cre; LRP1^loxP/loxP^* mice, but not in the liver compared with *LRP1^loxP/loxP^* mice (Supplementary Figure 2A–C). Serum and brain Aβ_40_ levels tended to be decreased, while hepatic levels were significantly elevated in *Vgat-Cre; LRP1^loxP/loxP^* mice (Supplementary Figure 2D–F). The ratio of Aβ_42/40_was significantly higher in *Vgat-Cre; LRP1^loxP/loxP^* mice than *LRP1^loxP/loxP^* mice (Supplementary Figure 2G). Deletion of LRP1 in GABAergic neurons tended to decrease hippocampal PSEN1 levels while it had no effect on PSEN1 levels in cortex and hippocampal APP levels (Supplementary Figure 2H–I).

### 3.7. Deficiency of LRP1 in GABAergic neurons causes neurodegeneration

We performed histopathological evaluation of the hippocampus to determine whether deleting LRP1 in GABAergic neurons causes neurodegeneration. Mice lacking LRP1 in GABAergic neurons showed necrotic changes such as neuronal shrinkage, darker cytoplasm and pyknosis in the hippocampus where mild microglial cell proliferation was also observed (Figure 6A). In addition, *Vgat-Cre; LRP1^loxP/loxP^* mice had satellite microglial cells surrounding some neurons (called perineuronal satellitosis) in the hippocampus, whereas the control mice showed normal cell morphology without satellitosis (Figure 6B). The quantifications of necrotic and glial cells and cells with satellitosis from the pathological images are shown in graphs (Figure 6C). To elucidate whether neurodegenerative changes are due to the LRP1 deletion or the impact of the obese phenotype, we performed the same histopathological evaluations before the mice became obese. The results revealed that increases in necrotic and glial cells are due to LRP1 deletion regardless of obesity state; however, satellitosis was only observed in older, obese mice (Figure 6D–F).

**Figure 6:**
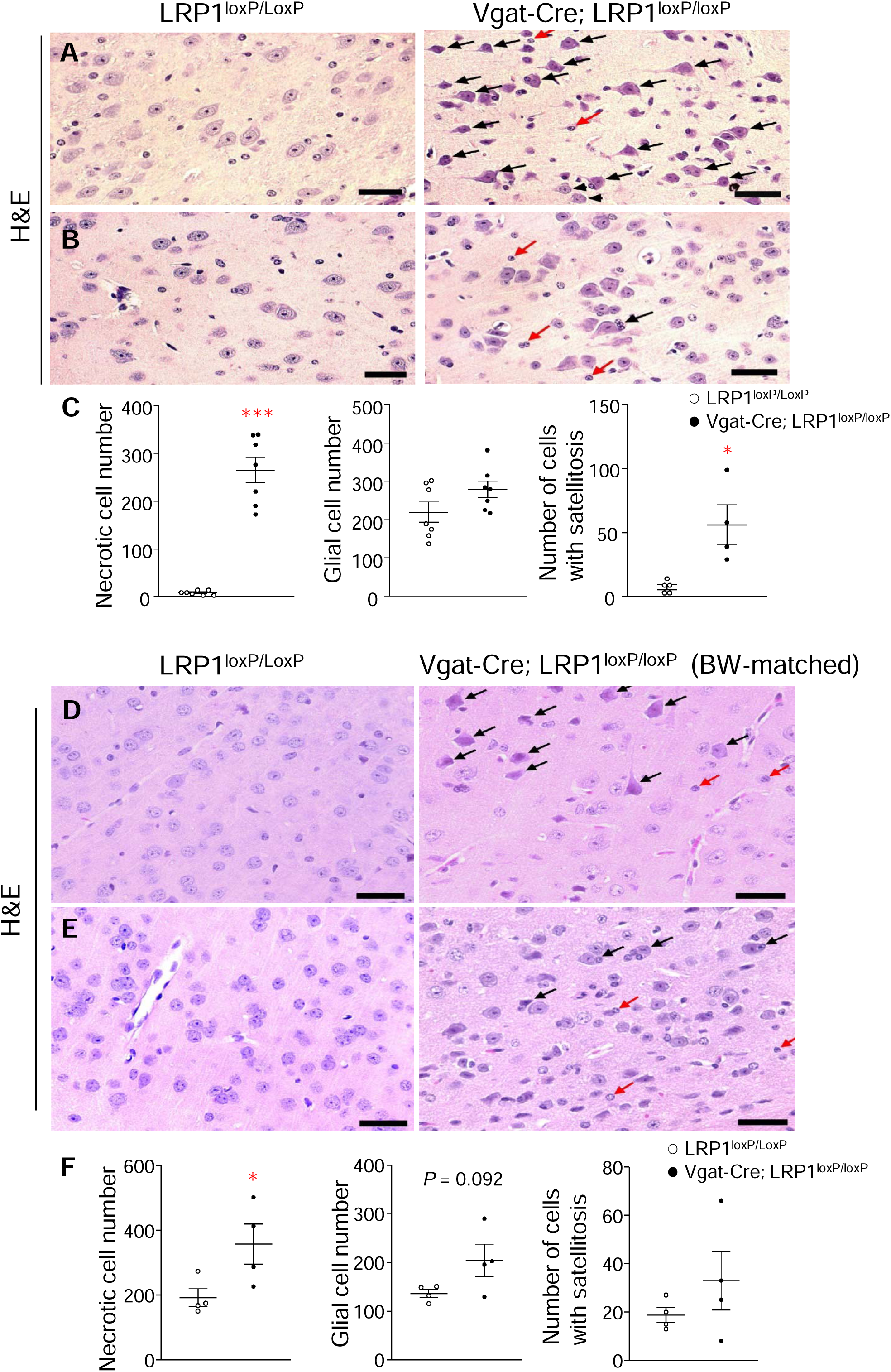
Histopathological features in the hippocampus of GABAergic neuron-specific LRP1-deficient mice before and after obesity. (**A**) Black arrows indicate necrotic changes in neurons and red arrows indicate glial cells in obese *Vgat-Cre; LRP1^loxP/loxP^* mice. Normal neuronal appearance of *LRP1^loxP/LoxP^* mice was seen (*n* = 7/group). (**B**) Black arrows indicates neurons with satellitosis and red arrows indicate glial cells in the hippocampus of obese *Vgat-Cre; LRP1^loxP/loxP^* mice. No satelliotosis in *LRP1^loxP/LoxP^* mice were found (*n* = 5/group). (**C**) The quantifications of neurodegenerative changes after obesity. (**D**) Black arrows show necrotic changes in neurons and red arrows show glial cells in normal *Vgat-Cre; LRP1^loxP/loxP^* mice. Normal neuronal appearance of *LRP1^loxP/LoxP^* mice was seen (*n* = 4/group). (**E**) Black arrows show neurons with satellitosis and red arrows show glial cells in the section from hippocampus of normal *Vgat-Cre; LRP1^loxP/loxP^* mice. No stelliotosis in *LRP1^loxP/LoxP^* mice were found (*n* = 4/group). (**F**) The quantifications of neurodegenerative changes before obesity. Necrotic neurons and perineural satellitosis as well as glial cells in the hippocampus were counted manually from five serial sections before and after obesity. Graphs represent the quantitation of necrotic neuron number, glial cell number, and number of neurons with satellitosis in 15 areas of each section. Scale bars represent 30 μm. All graphs show means ± SEM. \**P* <0.05, \*\*\**P* <0.001 vs. *LRP1^loxP/loxP^* by two-sided Student’s t-test or Mann-Whitney U test. H&E: Hematoxylin and eosin.

We further performed IHC for GFAP, IBA1, GAD67, and NMDAR in the hippocampus of *Vgat-Cre; LRP1^loxP/loxP^* mice. The number of immunoreactive GFAP^+^, IBA1^+^ and NMDAR^+^ neurons in hippocampus were significantly higher in *Vgat-Cre; LRP1^loxP/loxP^* mice compared to *LRP1^loxP/loxP^* mice (Figure 7A–C). However, *Vgat-Cre; LRP1^loxP/loxP^* mice had fewer GAD67^+^ immunoreactive cells compared to *LRP1^loxP/loxP^* mice (Figure 7D). The quantitative data from the IHC images are shown in the graphs in Figure 7. Similar to the histological evaluation, we used the mice before they became obese to determine whether the immunohistochemical changes are due to LRP1 deletion or obesity. We found a reduction in GAD67 and increase in IBA-1 in the younger, non-obese LRP1-deleted mice, without significant changes in NMDAR and GFAP, which were exclusive to the obese phenotype (Figure 8). Collectively, these findings demonstrate that LRP1 deletion from GABAergic neurons causes neurodegeneration and significant histological changes that may contribute to memory dysfunction and neurodegenerative processes.

**Figure 7:**
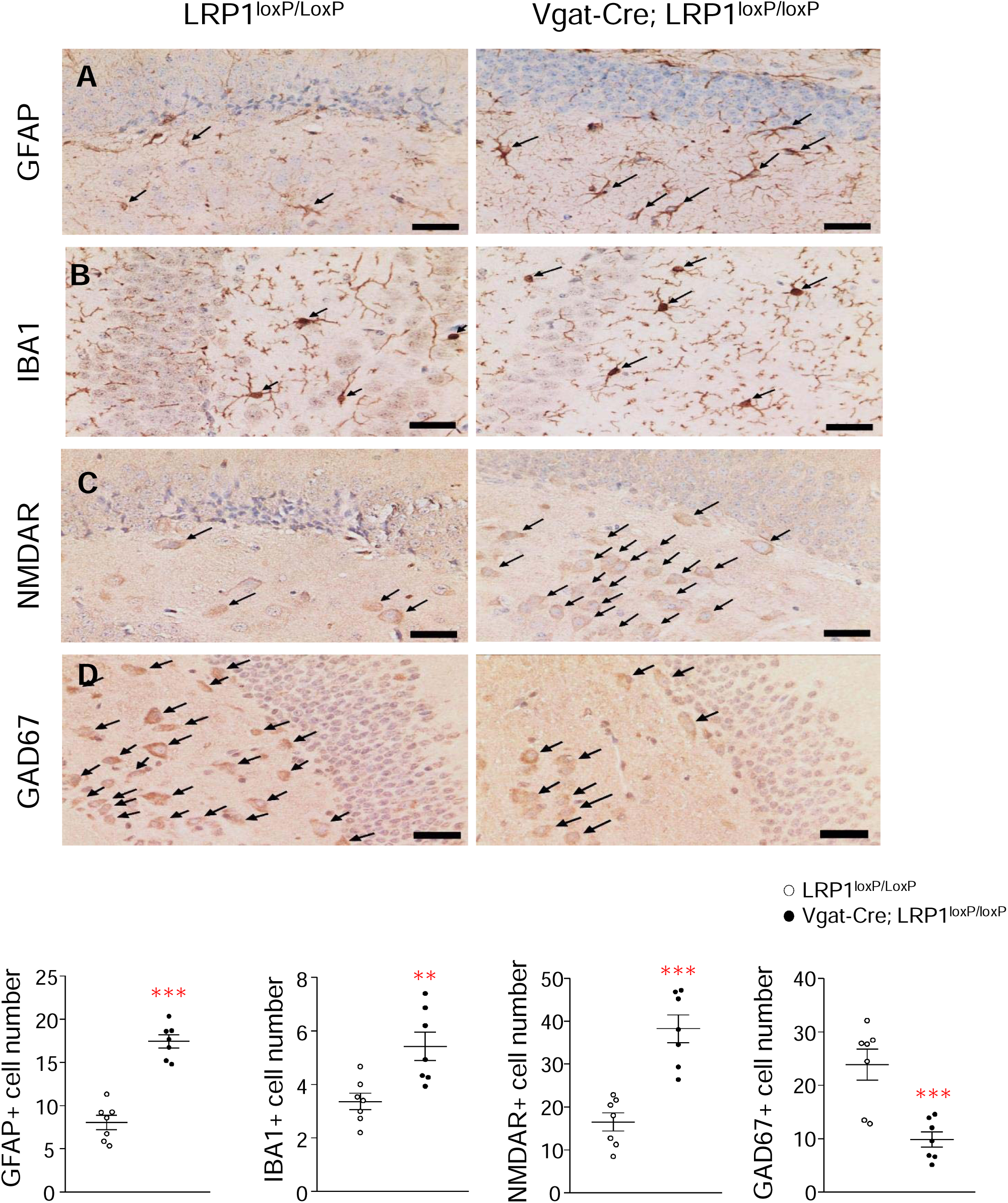
Immunohistochemical analysis in the hippocampus of GABAergic neuron-specific LRP1-deficient mice. (**A**) GFAP, (**B**) IBA1, (**C**) NMDAR, and (**D**) GAD67 immunoreactivity of *LRP1^loxP/loxP^* (*n* = 7) and *Vgat-Cre; LRP1^loxP/loxP^* (*n* = 7) male mice. Arrows indicate GFAP^+^, IBA1^+^, NMDAR^+^, and GAD67^+^ cells which were counted manually as described in Figure 6. Graphs represent the mean of quantitation of GFAP^+^, IBA1^+^, NMDAR^+^, and GAD67^+^ cells in 15 areas of each section. Scale bars represent 30 μm. All graphs show means ± SEM. \*\**P* <0.01, \*\*\**P* <0.001 vs. *LRP1^loxP/loxP^* by two-sided Student’s t-test.

**Figure 8:**
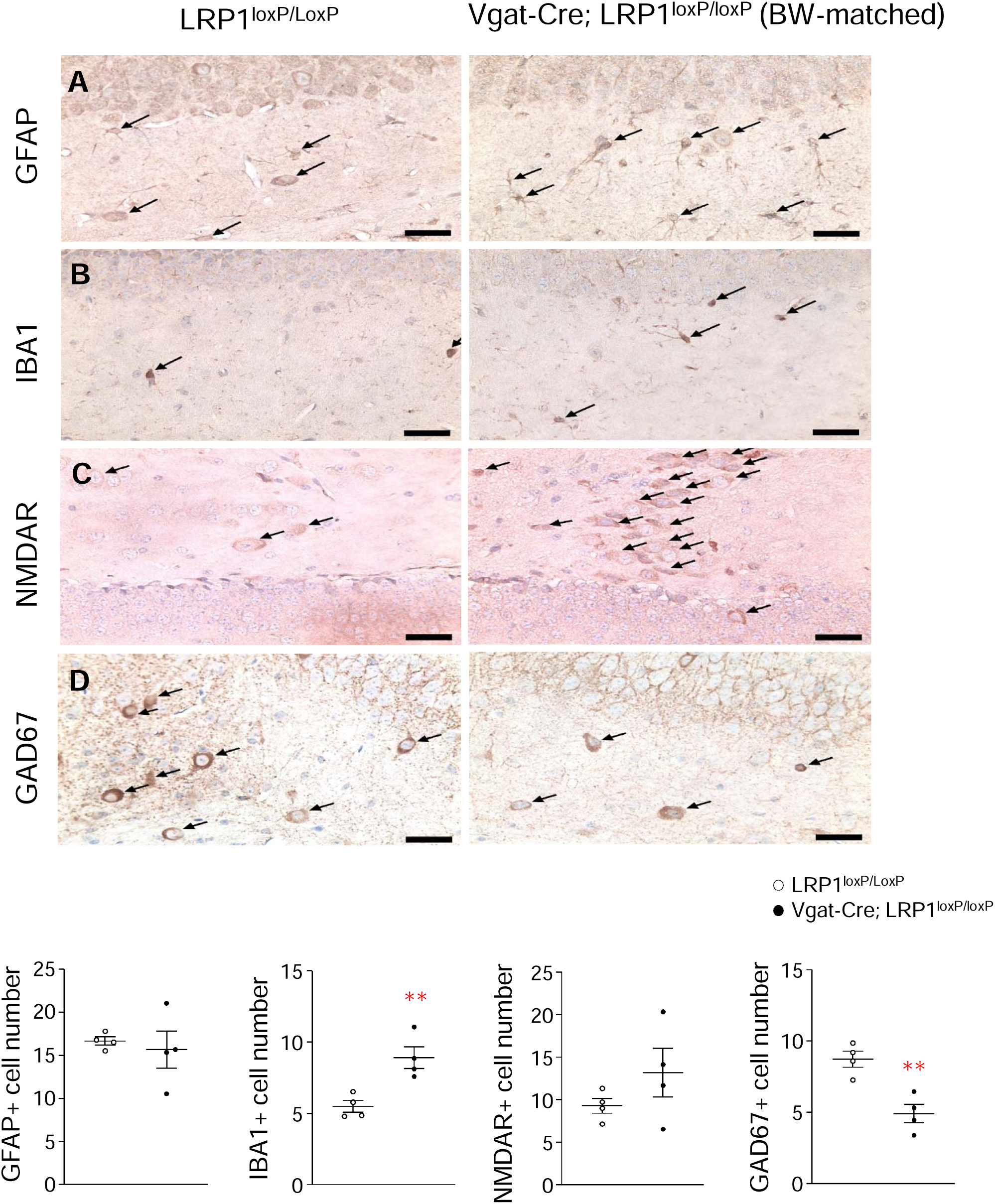
Immunohistochemical analysis in the hippocampus of GABAergic neuron-specific LRP1-deficient BW-matched mice. (**A**) GFAP, (**B**) IBA1, (**C**) NMDAR, and (**D**) GAD67 immunoreactivity of *LRP1^loxP/loxP^* (*n* = 4) and *Vgat-Cre; LRP1^loxP/loxP^* (*n* = 4) male mice. Arrows show GFAP^+^, IBA1^+^, NMDAR^+^, and GAD67^+^ cells which were counted manually as described in Figure 6. Graphs represent the mean of quantitation of GFAP^+^, IBA1^+^, NMDAR^+^, and GAD67^+^ cells in 15 areas of each section. Scale bars represent 30 μm. All graphs show means ± SEM. \*\**P* <0.01, \*\*\**P* <0.001 vs. *LRP1^loxP/loxP^* by two-sided Student’s t-test.

## 4. DISCUSSION

We showed that LRP1 deficiency in GABAergic neurons, which is known to cause obesity and metabolic disturbances, alters locomotor activity and impairs short-and long-term memory. Changes in obesity-related metabolic parameters caused by LRP1 deletion in GABAergic neurons correlate with declining cognitive performance. Pathological and IHC analysis demonstrate that LRP1 in GABAergic neurons plays a pivotal role in the progression of neuronal degeneration. Thus, LRP1 in GABAergic neurons is an important regulator of memory and cognitive function.

Our study revealed that the specific deletion of LRP1 from GABAergic neurons led to moderate impairment of motor coordination and locomotor activity. This is consistent with previous studies using general deletion of LRP1 in the forebrain of mice, suggesting an important role for LRP1 in GABAergic neurons in motor function [36,37]. However, previous studies demonstrated that reduction in motor coordination is also accompanied by increased adiposity, increased circulating insulin and leptin levels, glucose intolerance [38,39], and crosstalk between leptin and insulin signaling [40-42]. Moreover, while GABAergic LRP1 deleted mice were hypoactive and obese, non-obese mice with pan-neuronal LRP1 deletion exhibited hyperactive characteristics [37]. Thus, it is likely that neuronal dysfunction caused by GABAergic LRP1 deletion plays a role in the development of motor abnormalities, along with impaired energy balance.

Obesity is associated with memory and learning deficits [43,44]. One potential mechanism behind this suggests that excessive insulin causes an increased cerebral inflammatory response as well as increased Aβ levels, which may contribute to the deleterious effects of obesity on working memory [45]. Supporting this, our memory behavior tests (water T-maze and fear conditioning) and correlation analysis collectively confirmed that obesity and obesity-associated metabolic changes such as hyperinsulinemia, hyperleptinemia and high blood levels of ApoJ are correlated with poor memory performance. Moreover, histopathological and immunohistochemical analyses showed increased gliosis and neuronal death in obese GABAergic LRP1-deficient mice. However, non-obese mice with LRP1 deficiency in GABAergic cells also showed increase microglial proliferation and necrosis, which suggests that these histopathological findings are independent of obesity. In fact, transgenic mice for Alzheimer’s disease fed sucrose-sweetened water developed glucose intolerance and hyperinsulinemia in the absence of obesity, which exacerbated memory impairment and increased insoluble Aβ protein deposition [46]. Additionally, leptin has been shown to facilitate spatial learning and memory performance, and leptin deficiency led to impaired spatial memory in rodents [47-49]. In contrast, our LRP1-deficient mice with memory and learning deficits exhibited increased levels of leptin. Given that GABAergic LRP1-deficient mice are obese but not leptin resistant [14], it is conceivable that leptin action in our animals may not be involved in learning and memory performance.

The fact that a lack of LRP1 in GABAergic neurons led to cognitive dysfunction may be associated with the neurodegeneration observed in the hippocampus. The majority (80-90%) of the hippocampal neurons are principal cells, while the minority are GABAergic neurons (10-15%). However, the inhibitory GABAergic neurons are highly integrated and show functionally diverse properties, allowing them to control the spatio-temporal functions of the rest of the hippocampus [50,51]. Thus, physiological and morphological changes in these neurons may affect overall hippocampal function. The present findings indicate that a lack of LRP1 in GABAergic neurons causes moderate-to-severe neurodegeneration as evidenced by the increased number of glial cells and cells with necrosis and satellitosis in the hippocampal area. Histopathological and immunohistochemical evaluations raised the question as to whether these cells are GABAergic neurons. Morphophysiologically, excitatory principal cells of the hippocampus are pyramidal neurons located in specific areas such as the stratum pyramidale while GABAergic interneurons are scattered throughout the hippocampus [52]. We found a significant decrease in the number of inhibitory GAD67+ cells in both pre-obese and obese mice, while the number of glutamate responsive NMDAR+ neurons were unchanged or increased, respectively. Thus, the neurodegeneration in the hippocampus possibly involved the GABAergic neurons due to the deletion of LRP1 with a compensatory increase in NMDA responsive neurons as a consequence of obesity and/or LRP1 deletion. NMDAR-mediated excitatory glutamatergic neurotransmission critically regulate neuronal function, connectivity, and cell survival [53-56]. In the hippocampus, the cellular processes that control learning and memory are dependent on NMDAR [54,56]. On the other hand, excessive neuronal activity via NMDAR results in hyper-excitation and ultimately can cause excitotoxicity, neurodegeneration and cell death [57,58]. This may underlie the mechanism of obesity-associated cognitive impairment occurred in *Vgat-Cre; LRP1^loxP/loxP^* mice. Taken together, the cognitive dysfunction found in *Vgat-Cre; LRP1^loxP/loxP^* mice is likely due to the neurodegenerative changes in the hippocampus.

LRP1 in the brain also plays a crucial role in regulating Aβ production, as evidenced by the fact that a reduction in LRP1 expression is associated with Aβ clearence deficit and accumulation [19-22]. While Aβ_40_ accounts for the most of the Aβ production, Aβ_42_ is thought to trigger Aβ accumulation [59]. Therefore, changes in these two Aβ products provide critical information for Aβ accumulation. Even though inconsistencies have been reported [60], an increase in the Aβ_42/40_ ratio is considered to be a surogate for the Aβ accumulation in Alzheimer’s disease [61]. In this respect, our data show that mice lacking GABAergic LRP1 have increased serum and brain Aβ_42/40_ ratios. Although there is no evidence supporting a role of naturally occuring Aβ_42_ in cognitive imparement in mice, this may have important consequences in humans and future transgenic AD mice studies. Thus, Aβ accumulation could be a factor for the progression of cognitive decline in AD models lacking LRP1 in GABAergic neurons. In addition, other than the direct effect of LRP1 deletion, the low locomotor activity of the LRP1 knockout mice due to obesity might have further contributed to the Aβ accumulation as physical activity affects Aβ metabolism [62-64]. The findings indicate that LRP1 expressions are changed in other cells as a compensatory mechanism. PSEN1 expression was also slightly down-regulated in response to LRP1 deletion from GABAergic neurons. However, the overall impact might not have significant effects on the dynamics of Aβ because the compensatory changes are not substantial even though statistically significant. Of note, the change in Aβ_40_ in the liver might be due to the metabolic impact of obesity and not a result of GABAergic LRP1 knowdown [65].

## 5. CONCLUSIONS

LRP1 deletion in GABAergic neurons resulted in behavior abnormalities, and memory and learning deficits. These effects were accompanied by neurodegeneration, neuroinflammation, which preceded the onset of obesity. Thus, LRP1 in GABAergic neurons may be an important regulator of memory and cognitive function.

## AUTHOR CONTRIBUTION

**Aaron Aykut Uner, Zhi-Shuai Hou, Woojin S. Kim, Vincent Prevot, Barbara J. Caldarone, Hyon Lee and Young-Bum Kim:** Designed and planned the study. **Aaron Aykut Uner, Zhi-Shuai Hou and Barbara J. Caldarone:** Performed most of behavioral experiments. **Aaron Aykut Uner and Ahmet Aydogan:** Performed histopathology and IHC experiments. **Kellen C.C. Rodrigues, Jennie Young, Anthony Choi and Won-Mo Yang:** Carried out blood serum parameters and immunoblotting analysis. **Bradley T. Hyman:** Conceptualized the study and interpreted the data. **Aaron Aykut Uner, Zhi-Shuai Hou, Ahmet Aydogan, Hyon Lee and Young-Bum Kim:** Wrote the manuscript with input from all other authors.

## DATA AVAILABILITY

The authors confirm that the data are available from the corresponding author upon request.

### Abbreviations

LRP1: low-density lipoprotein receptor-related protein-1;
LDL: low-density lipoprotein;
Aβ: amyloid beta;
HMS: Harvard Medical School;
Vgat: vesicular GABA transporter;
ELISA: enzyme linked immunosorbent assay;
HOMA-IR: homeostatic model assessment for insulin resistance;
PSEN1: presenilin-1;
APP: amyloid precursor protein;
TBS: tris-buffered saline;
H&E: hematoxylin and eosin;
IHC: Immunohistochemistry;
ABC: avidin–biotin–peroxidase complex;
GFAP: glial fibrillary acidic protein;
IBA1: ionized calcium binding adaptor molecule 1;
GAD67: glutamic acid decarboxylase 67;
DAB-H2O2: 3,3’-diaminobenzidine tetrahydrochloride;
AF: Alexa fluor;
FFA: free fatty acids;
ApoJ: apolipoprotein J;
CCFC: contextual and cued fear conditioning tasks;
NMDARs: N-methyl-d-aspartate receptors;
GABA: gamma-aminobutyric acid;
PBS: phosphate-buffered saline;
ANOVA: analysis of variance

## ACKNOWLEDGMENTS

This work was supported by grants from the National Institutes of Health (3R01DK106076-03S1, 3R01 DK123002-02S1, R01DK106076-04S1, R01AG080842 to YBK). A.A. is a recipient of the fellowship from the Scientific and Technological Research Council of Turkey (TUBITAK, 1059B192100216) and K.C.R is a recipient of the fellowship from the São Paulo Research Foundation from Brazil (FAPESP 2019/19938-5). We thank Brad Lowell for the *Vgat-IRES-Cre* recombinase mice, and all members of the Kim lab for their helpful advice and discussions. We also thank Dr. Jongkyun Kang (Harvard Medical School, Boston MA) for his help with the interpretation of the behavioral data. We are grateful to the Histology Core at Beth Israel Deaconess Medical Center.

## REFERENCES

1. Anstey, K.J., Cherbuin, N., Budge, M., Young, J., 2011. Body mass index in midlife and late-life as a risk factor for dementia: a meta-analysis of prospective studies. Obes Rev 12:e426–437.

2. Fitzpatrick, A.L., Kuller, L.H., Lopez, O.L., Diehr, P., O’Meara, E.S., Longstreth Jr, W.T., et al. 2009. Midlife and late-life obesity and the risk of dementia: cardiovascular health study. Arch Neurol 66:336–342.

3. Jahn, H., 2013. Memory loss in Alzheimer’s disease. Dialogues Clin Neurosci 15:445–454.

4. Lane, C.A., Hardy, J., Schott, J.M., 2018. Alzheimer’s disease. Eur J Neurol 25:59–70.

5. Profenno, L.A., Porsteinsson, A.P., Faraone, S.V., 2010. Meta-analysis of Alzheimer’s disease risk with obesity, diabetes, and related disorders. Biol Psychiatry 67:505–512.

6. Serlin, Y., Levy, J., Shalev, H., 2011. Vascular pathology and blood-brain barrier disruption in cognitive and psychiatric complications of type 2 diabetes mellitus. Cardiovasc Psychiatry Neurol 609202.

7. Cho, S., Lee, H., Seo, J., 2021. Impact of Genetic Risk Factors for Alzheimer’s Disease on Brain Glucose Metabolism. Mol Neurobiol 58:2608–2619.

8. de Leon, M.J., Convit, A., Wolf, O.T., Tarshish, C.Y., DeSanti, S., Rusinek, H., et al. 2001. Prediction of cognitive decline in normal elderly subjects with 2-[(18)F]fluoro-2-deoxy-D-glucose/poitron-emission tomography (FDG/PET). Proc Natl Acad Sci U S A 98:10966–10971.

9. Drzezga, A., Lautenschlager, N., Siebner, H., Riemenschneider, M., Willoch, F., Minoshima, S., et al. 2003. Cerebral metabolic changes accompanying conversion of mild cognitive impairment into Alzheimer’s disease: a PET follow-up study. Eur J Nucl Med Mol Imaging 30:1104–1113.

10. Gordon, B.A., Blazey, T.M., Su, Y., Hari-Raj, A., Dincer, A., Flores, S., et al. 2018. Spatial patterns of neuroimaging biomarker change in individuals from families with autosomal dominant Alzheimer’s disease: a longitudinal study. Lancet Neurol 17:241–250.

11. Liu, F., Shi, J., Tanimukai, H., Gu, J., Gu, J., Grundke-Iqbal, I., et al. 2009. Reduced O-GlcNAcylation links lower brain glucose metabolism and tau pathology in Alzheimer’s disease. Brain 132:1820–1832.

12. Moloney, A.M., Griffin, R.J., Timmons, S., O’Connor, R., Ravid, R., O’Neill, C., 2010. Defects in IGF-1 receptor, insulin receptor and IRS-1/2 in Alzheimer’s disease indicate possible resistance to IGF-1 and insulin signalling. Neurobiol Aging 31:224-243.

13. Mosconi, L., Tsui, W.H., Rusinek, H., De Santi, S., Li, Y., Wang, G.J., et al. 2007. Quantitation, regional vulnerability, and kinetic modeling of brain glucose metabolism in mild Alzheimer’s disease. Eur J Nucl Med Mol Imaging 34:1467–1479.

14. Kang, M.C., Seo, J.A., Lee, H., Uner, A., Yang, W.M., Rodrigues, K., et al. 2021. LRP1 regulates food intake and energy balance in GABAergic neurons independently of leptin action. Am J Physiol Endocrinol Metab 320:E379–E389.

15. Liu, Q., Zhang, J., Zerbinatti, C., Zhan, Y., Kolber, B.J., Herz, J., et al. 2011. Lipoprotein receptor LRP1 regulates leptin signaling and energy homeostasis in the adult central nervous system. PLoS Biol 9:e1000575.

16. Fuentealba, R.A., Liu, Q., Kanekiyo, T., Zhang, J., Bu, G., 2009. Low density lipoprotein receptor-related protein 1 promotes anti-apoptotic signaling in neurons by activating Akt survival pathway. J Biol Chem 284:34045–34053.

17. Bu, G., 2009. Apolipoprotein E and its receptors in Alzheimer’s disease: pathways, pathogenesis and therapy. Nat Rev Neurosci 10:333–344.

18. Lillis, A.P., Van Duyn, L.B., Murphy-Ullrich, J.E., Strickland, D.K., 2008. LDL receptor-related protein 1: unique tissue-specific functions revealed by selective gene knockout studies. Physiol Rev 88:887–918.

19. Shibata, M., Yamada, S., Kumar, S.R., Calero, M., Bading, J., Frangione, B., et al. 2000. Clearance of Alzheimer’s amyloid-ss(1-40) peptide from brain by LDL receptor-related protein-1 at the blood-brain barrier. J Clin Invest 106:1489–1499.

20. Van Uden, E., Mallory, M., Veinbergs, I., Alford, M., Rockenstein, E., Masliah, E., 2002. Increased extracellular amyloid deposition and neurodegeneration in human amyloid precursor protein transgenic mice deficient in receptor-associated protein. J Neurosci 22:9298–9304.

21. Fuentealba, R.A., Liu, Q., Zhang, J., Kanekiyo, T., Hu, X., Lee, J.M., et al. 2010. Low-density lipoprotein receptor-related protein 1 (LRP1) mediates neuronal Abeta42 uptake and lysosomal trafficking. PLoS One 5:e11884.

22. Kang, D.E., Pietrzik, C.U., Baum, L., Chevallier, N., Merriam, D.E., Kounnas, M.Z., et al. 2000. Modulation of amyloid beta-protein clearance and Alzheimer’s disease susceptibility by the LDL receptor-related protein pathway. J Clin Invest 106:1159–1166.

23. Storck, S.E., Meister, S., Nahrath, J., Meißner, J.N., Schubert, N., Di Spiezio, A., et al. 2016. Endothelial LRP1 transports amyloid-beta(1-42) across the blood-brain barrier. J Clin Invest 126:123–136.

24. Knopp, J.L., Holder-Pearson, L., Chase, J.G., 2019. Insulin Units and Conversion Factors: A Story of Truth, Boots, and Faster Half-Truths. J Diabetes Sci Technol 13:597–600.

25. Matthews, D.R., Hosker, J.P., Rudenski, A.S., Naylor, B.A., Treacher, D.F., Turner, R.C., 1985. Homeostasis model assessment: insulin resistance and beta-cell function from fasting plasma glucose and insulin concentrations in man. Diabetologia 28:412–419.

26. Luna, L.G., 1968. Manual of histologic staining methods of the armed forces institute of pathology. 3rd ed. New York: McGraw-Hill.

27. Charnay, Y., Imhof, A., Vallet, P.G., Kovari, E., Bouras, C., Giannakopoulos, P., 2012. Clusterin in neurological disorders: molecular perspectives and clinical relevance. Brain Res Bull 88:434–443.

28. Elias-Sonnenschein, L.S., Bertram, L., Visser, P.J., 2012. Relationship between genetic risk factors and markers for Alzheimer’s disease pathology. Biomark Med 6:477–495.

29. Nuutinen, T., Suuronen, T., Kauppinen, A., Salminen, A., 2009. Clusterin: a forgotten player in Alzheimer’s disease. Brain Res Rev 61:89–104.

30. Yu, J.T., Tan, L., 2012. The role of clusterin in Alzheimer’s disease: pathways, pathogenesis, and therapy. Mol Neurobiol 45:314–326.

31. Hammad, S.M., Ranganathan, S., Loukinova, E., Twal, W.O., Argraves, W.S., 1997. Interaction of apolipoprotein J-amyloid beta-peptide complex with low density lipoprotein receptor-related protein-2/megalin. A mechanism to prevent pathological accumulation of amyloid beta-peptide. J Biol Chem 272:18644–18649.

32. Kounnas, M.Z., Moir, R.D., Rebeck, G.W., Bush, A.I., Argraves, W.S., Tanzi, R.E., et al. 1995. LDL receptor-related protein, a multifunctional ApoE receptor, binds secreted beta-amyloid precursor protein and mediates its degradation. Cell 82:331–340.

33. Kwon, M.J., Ju, T.J., Heo, J.Y., Kim, Y.W., Kim, J.Y., Won, K.C., et al. 2014. Deficiency of clusterin exacerbates high-fat diet-induced insulin resistance in male mice. Endocrinology 155:2089–2101.

34. Seo, J.A., Kang, M.C., Ciaraldi, T.P., Kim, S.S., Park, K.S., Choe, C., et al. 2018. Circulating ApoJ is closely associated with insulin resistance in human subjects. Metabolism 78:155–166.

35. Won, J.C., Park, C.Y., Oh, S.W., Lee, E.S., Youn, B.S., Kim, M.S., 2014. Plasma clusterin (ApoJ) levels are associated with adiposity and systemic inflammation. PLoS One 9:e103351.

36. Liu, Q., Trotter, J., Zhang, J., Peters, M.M., Cheng, H., Bao, J., et al. 2010. Neuronal LRP1 knockout in adult mice leads to impaired brain lipid metabolism and progressive, age-dependent synapse loss and neurodegeneration. J Neurosci 30:17068–17078.

37. May, P., Rohlmann, A., Bock, H.H., Zurhove, K., Marth, J.D., Schomburg, E.D., et al. 2004. Neuronal LRP1 functionally associates with postsynaptic proteins and is required for normal motor function in mice. Mol Cell Biol 24:8872–8883.

38. Stojakovic, A., Mastronardi, C.A., Licinio, J., Wong, M.L., 2019. Long-term consumption of high-fat diet impairs motor coordination without affecting the general motor activity. J Transl Sci 6:1–10.

39. Smith, S.M., Pjetri, E., Friday, W.B., Presswood, B.H., Ricketts, D.K., Walter, K.R., et al. 2022. Aging-Related Behavioral, Adiposity, and Glucose Impairments and Their Association following Prenatal Alcohol Exposure in the C57BL/6J Mouse. Nutrients 14:1438.

40. Huo, L., Gamber, K., Greeley, S., Silva, J., Huntoon, N., Leng, X.H., et al., 2009. Leptin-dependent control of glucose balance and locomotor activity by POMC neurons. Cell Metabolism 9:537–547.

41. Sartorius, T., Heni, M., Tschritter, O., Preissl, H., Hopp, S., Fritsche, A., et al. 2012. Leptin affects insulin action in astrocytes and impairs insulin-mediated physical activity. Cell Physiol Biochem 30:238–246.

42. Wang, C., Chan, J.S., Ren, L., Yan, J.H., 2016. Obesity Reduces Cognitive and Motor Functions across the Lifespan. Neural Plast 2016:2473081.

43. Coppin, G., Nolan-Poupart, S., Jones-Gotman, M., Small, D.M., 2014. Working memory and reward association learning impairments in obesity. Neuropsychologia 65:146–155.

44. Gunstad, J., Paul, R.H., Cohen, R.A., Tate, D.F., Gordon, E., 2006. Obesity is associated with memory deficits in young and middle-aged adults. Eat Weight Disord 11:e15–19.

45. Craft, S., 2005. Insulin resistance syndrome and Alzheimer’s disease: age-and obesity-related effects on memory, amyloid, and inflammation. Neurobiol Aging 26 Suppl 1:65–69.

46. Cao, D., Lu, H., Lewis, T.L., Li, L., 2007. Intake of sucrose-sweetened water induces insulin resistance and exacerbates memory deficits and amyloidosis in a transgenic mouse model of Alzheimer disease. J Biol Chem 282:36275–36282.

47. Li, X.L., Aou, S., Oomura, Y., Hori, N., Fukunaga, K., Hori, T., 2002. Impairment of long-term potentiation and spatial memory in leptin receptor-deficient rodents. Neuroscience 113:607–615.

48. Oomura, Y., Aou, S., Fukunaga, K., 2010. Prandial increase of leptin in the brain activates spatial learning and memory. Pathophysiology 17:119–127.

49. Oomura, Y., Hori, N., Shiraishi, T., Fukunaga, K., Takeda, H., Tsuji, M., et al. 2006. Leptin facilitates learning and memory performance and enhances hippocampal CA1 long-term potentiation and CaMK II phosphorylation in rats. Peptides 27:2738–2749.

50. Booker, S.A., Vida, I., 2018. Morphological diversity and connectivity of hippocampal interneurons. Cell Tissue Res 373:619–641.

51. Vida, I., Degro, C.E., Booker., S.A., 2018. Morphology of Hippocampal Neurons. In: Cobb, S., Cutsuridis, V., Graham, B.P., Vida, I., editors. Hippocampal Microcircuits: A Computational Modeler’s Resource Book, New York: Springer, p. 27–67.

52. Pelkey, K.A., Chittajallu, R., Craig, M.T., Tricoire, L., Wester, J.C., McBain, C.J., 2017. Hippocampal GABAergic Inhibitory Interneurons. Physiol Rev 97:1619–1747.

53. Brigman, J.L., Wright, T., Talani, G., Prasad-Mulcare, S., Jinde, S., Seabold, G.K., et al., 2010. Loss of GluN2B-containing NMDA receptors in CA1 hippocampus and cortex impairs long-term depression, reduces dendritic spine density, and disrupts learning. The Journal of Neuroscience 30:4590–4600.

54. Sakimura, K., Kutsuwada, T., Ito, I., Manabe, T., Takayama, C., Kushiya, E., et al., 1995. Reduced hippocampal LTP and spatial learning in mice lacking NMDA receptor epsilon 1 subunit. Nature 373:151–155.

55. Tolias, K.F., Bikoff, J.B., Burette, A., Paradis, S., Harrar, D., Tavazoie, S., et al., 2005. The Rac1-GEF Tiam1 couples the NMDA receptor to the activity-dependent development of dendritic arbors and spines. Neuron 45:525–538.

56. Wong, R.O., Ghosh, A., 2002. Activity-dependent regulation of dendritic growth and patterning. Nat Rev Neurosci 3:803–812.

57. Choi, D.W., 1992. Excitotoxic cell death. J Neurobiol 23:1261–1276.

58. Tymianski, M., Charlton, M.P., Carlen, P.L., Tator, C.H., 1993. Source specificity of early calcium neurotoxicity in cultured embryonic spinal neurons. J Neurosci 13:2085–2104.

59. Kelleher, R.J. 3rd., Shen, J., 2017. Presenilin-1 mutations and Alzheimer’s disease. Proc Natl Acad Sci U S A 114:629-631.

60. Sun, L., Zhou, R., Yang, G., Shi, Y., 2017. Analysis of 138 pathogenic mutations in presenilin-1 on the in vitro production of Abeta42 and Abeta40 peptides by gamma-secretase. Proc Natl Acad Sci U S A 114:E476–E485.

61. Selkoe, D.J., Hardy, J., 2016. The amyloid hypothesis of Alzheimer’s disease at 25 years. EMBO Mol Med 8:595–608.

62. Lonnemann, N., Korte, M., Hosseini, S., 2023. Repeated performance of spatial memory tasks ameliorates cognitive decline in APP/PS1 mice. Behav Brain Res 438:114218.

63. Maliszewska-Cyna, E., Vecchio, L.M., Thomason, L.A.M., Oore, J.J., Steinman, J., Joo, I.L., et al. 2020. The effects of voluntary running on cerebrovascular morphology and spatial short-term memory in a mouse model of amyloidosis. Neuroimage 222:117269.

64. Rangasamy, S.B., Jana, M., Dasarathi, S., Kundu, M., Pahan, K., 2023. Treadmill workout activates PPARalpha in the hippocampus to upregulate ADAM10, decrease plaques and improve cognitive functions in 5XFAD mouse model of Alzheimer’s disease. Brain Behav Immun 109:204–218.

65. Bosoi, C.R., Vandal, M., Tournissac, M., Leclerc, M., Fanet, H., Mitchell, P.L., et al. 2021. High-Fat Diet Modulates Hepatic Amyloid beta and Cerebrosterol Metabolism in the Triple Transgenic Mouse Model of Alzheimer’s Disease. Hepatol Commun 5:446–460.

