## Supplementary Figure 1 and 2 for "GABAergic LRP1 is a key link between obesity and memory function"

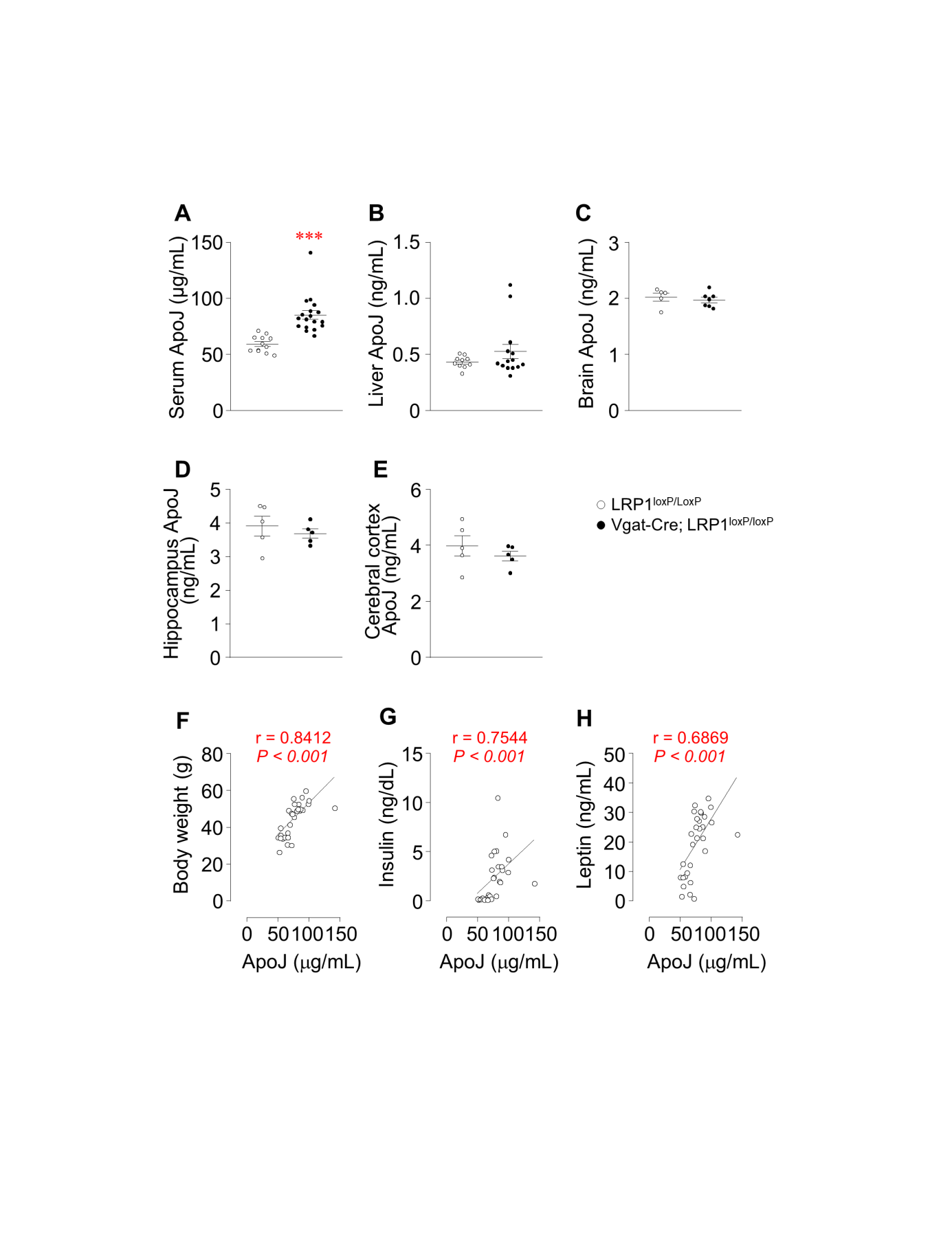

**Supplementary Figure 1:** **LRP1 deficiency in GABAergic neurons increases serum ApoJ levels.** (**A**) Serum ApoJ (*n* = 12 for control, *n* = 18 for *Vgat-Cre; LRP1^loxP/loxP^*), (**B**) liver ApoJ (*n* = 10 for control, *n* = 14 for *Vgat-Cre; LRP1^loxP/loxP^*), (**C**) brain ApoJ (*n* = 5 for control, *n* = 7 for *Vgat-Cre; LRP1^loxP/loxP^*), (**D**) hippocampus ApoJ (*n* = 5 for control, *n* = 5 for *Vgat-Cre; LRP1^loxP/loxP^*), and (**E**) cerebral cortex ApoJ (*n* = 5 for control, *n* = 5 for *Vgat-Cre; LRP1^loxP/loxP^*) of *LRP1^loxP/loxP^* and *Vgat-Cre; LRP1^loxP/loxP^* male mice. Relationship of serum ApoJ with (**F**) body weight, (**G**) serum insulin, and (**H**) serum leptin in *LRP1^loxP/loxP^* (*n* = 12) and *Vgat-Cre; LRP1^loxP/loxP^* (*n* = 18) male mice. ApoJ levels of serum and tissues were measured from overnight fasted mice at 32 weeks of age. The *P* values were obtained by Spearman correlation analysis and r values indicate Spearman correlation coefficient. All graph show means ± SEM. ****P* <0.001 vs. *LRP1^loxP/loxP^* by two-sided Student’s t-test.

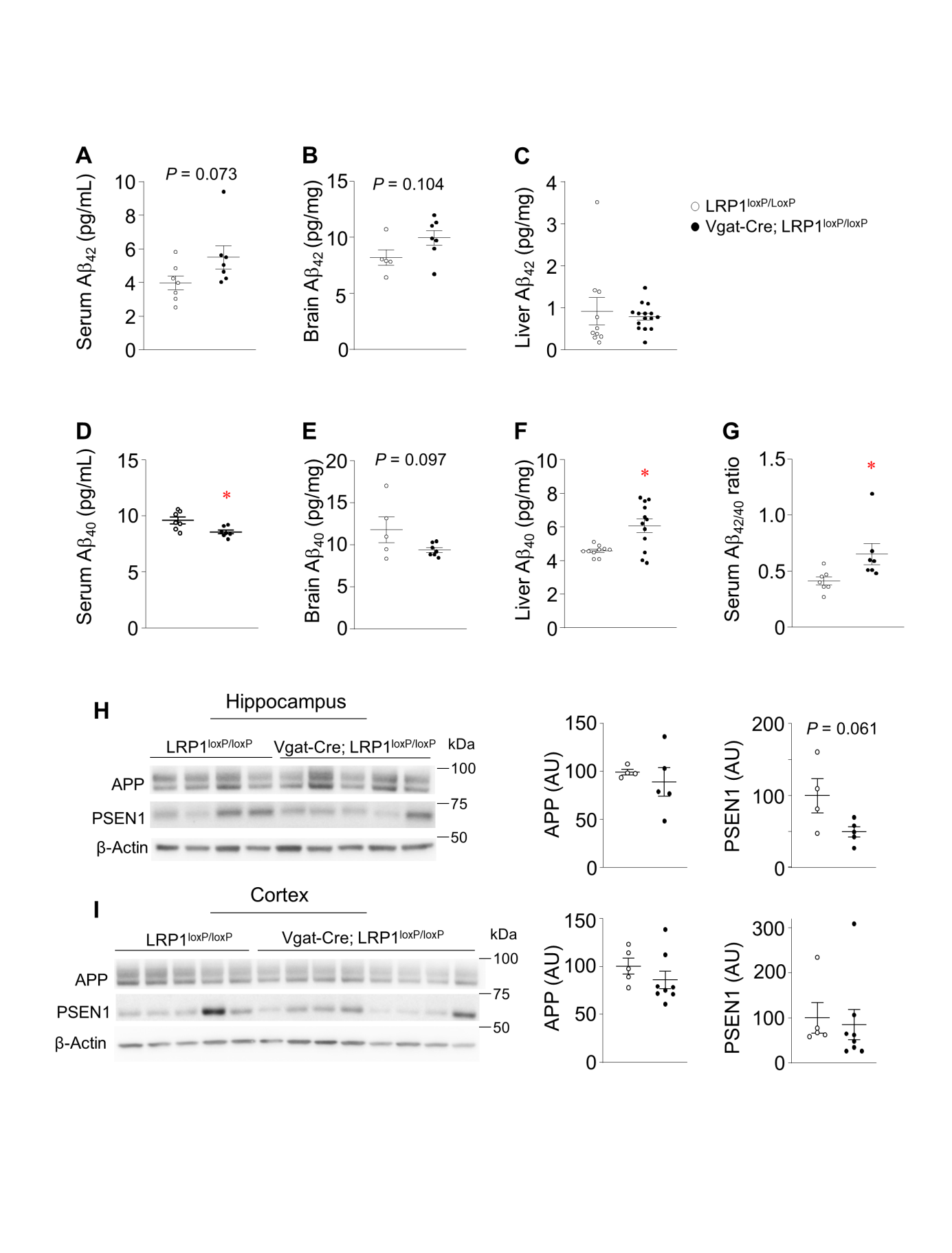
**Supplementary Figure 2:** **The impact of LRP1 on amyloid-β accumulation.** (**A**) Serum Aβ_42_ level (*n* = 7 for control, *n* = 7 for *Vgat-Cre; LRP1^loxP/loxP^*), (**B**) brain Aβ_42_ level (*n* = 5 for control, *n* = 7 for *Vgat-Cre; LRP1^loxP/loxP^*), (**C**) liver Aβ_42_ level (*n* = 10 for control, *n* = 15 for *Vgat-Cre; LRP1^loxP/loxP^*), (**D**) serum Aβ_40_ level (*n* = 7 for control, *n* = 7 for *Vgat-Cre; LRP1^loxP/loxP^*) (**E**) brain Aβ_40_ level (*n* = 5 for control, *n* = 7 for *Vgat-Cre; LRP1^loxP/loxP^*), (**F**) liver Aβ_40_ level (*n* = 10 for control, *n* = 12 for *Vgat-Cre; LRP1^loxP/loxP^*), and (**G**) serum Aβ_42/40_ ratio of male *LRP1^loxP/loxP^* and *Vgat-Cre; LRP1^loxP/loxP^* male mice (32 weeks old). APP and PSEN1 protein levels in (**H**) hippocampus and (**I**) cortex of *LRP1^loxP/loxP^* and *Vgat-Cre; LRP1^loxP/loxP^* mice. Aβ_42_ and Aβ_40_ levels in serum and tissues were measured from overnight fasted mice at 32 weeks of age. All graphs show means ± SEM. **P* <0.05 vs. *LRP1^loxP/loxP^* by two-sided Student’s t-test or Mann-Whitney U test.
